# Investigation of reward learning and feedback sensitivity in non-clinical participants with a history of early life stress

**DOI:** 10.1101/2020.11.13.380444

**Authors:** Matthew P Wilkinson, Chloe L Slaney, Jack R Mellor, Emma S J Robinson

## Abstract

Early life stress (ELS) is an important risk factor for the development of depression. Impairments in reward learning and feedback sensitivity are suggested to be an intermediate phenotype in depression aetiology therefore we hypothesised that similar impairments are present in healthy adults with a history of ELS. We recruited 64 adults with high levels of ELS and no diagnosis of a current mental health disorder and 65 controls. Participants completed the probabilistic reversal learning task and probabilistic reward task followed by depression, anhedonia, social status, and stress scales. Participants with high levels of ELS showed decreased positive feedback sensitivity in the probabilistic reversal learning task compared to controls. High ELS participants also trended towards possessing a decreased model-free learning rate. This was coupled with a decreased learning ability in the acquisition phase of block 1 following the practice session. Neither group showed a reward induced response bias in the probabilistic reward task however high ELS participants exhibited decreased stimuli discrimination. Due to the PRT not meeting its primary endpoint a separate cohort of control participants were tested in a modified PRT where they showed a response bias. This indicates the PRT can be successfully carried out online. Overall, these data suggest that healthy participants without a mental health diagnosis and high levels of ELS show deficits in positive feedback sensitivity and reward learning in the probabilistic reversal learning task that are distinct from depressed patients. These deficits may be relevant to increased depression vulnerability.

## 1. Introduction

Early life stress (ELS) is a major known risk factor for the development of depression [1–4]. ELS has also been found to lower the threshold of stress required to precipitate depression [5], one of the major triggers in healthy populations [6]. Elevated levels of childhood stress lead to widespread functional and morphological alterations in the adult brain with the hippocampus, amygdala and prefrontal cortex being most impacted [7,8]. Amongst other functions, these regions are vital mediators of reward learning: the ability of reward to modulate future behaviour [9–16]. However, how ELS influences the developing brain to predispose individuals to psychiatric illness is not yet understood.

Reward learning deficits have been proposed to be an intermediate phenotype in the aetiology and maintenance of depression [17–20]. Depressed patients show decreased reward sensitivity in the probabilistic reward task (PRT), a test of reward learning [19]. These deficits have been observed to both predict the risk of disease development [21] and persistence [18,22]. Utilising a different reward learning assay, the probabilistic reversal learning task (PRLT), depressed patients show impaired accuracy following probabilistic rule reversal and increased sensitivity to probabilistic negative feedback [23,24]. Acute stress has also been observed to impair reward learning [25,26] suggesting a potential link between stress, reward processing deficits and depression aetiology. Therefore this suggests that a divergent developmental trajectory in regions involved with reward processing as seen in ELS could lead to depression vulnerability through impaired reward learning as an intermediate phenotype.

Previous studies have therefore investigated reward processing deficits in people who have experienced ELS. Hanson and colleagues [27] recruited adolescents with a history of physical abuse who then completed a probabilistic learning task where they showed lower associative learning compared to controls. Changes in reward learning have also been reported within another probabilistic reward task, the probabilistic stimulus selection task (PSST). Women with a history of childhood sexual abuse and a diagnosis of Major Depressive disorder (MDD) showed decreased performance on trials requiring learning of previously rewarded information compared to MDD only and control groups [28]. Although these studies provide valuable insights, they use different tasks to those previously used to study depressed populations making direct comparisons difficult. Additionally, studies are needed in adults without a current mental health diagnosis to understand if any reward processing changes are present in individuals at higher risk of mental health disorders.

In this study it was hypothesised that ELS is associated with alterations in reward processing and feedback sensitivity in an otherwise healthy adult population. Two groups of adult participants that self-reported no current diagnosis of a mental health condition or Parkinson’s disease were recruited online and completed a survey of adverse childhood experiences [29] before being split into high and no ELS groups based upon this. Participants completed the PRT (points based) and PRLT with PRLT data additionally being analysed using a Q-learning model to probe reward learning parameter changes. Participants were asked about stress exposure to enable exploratory analysis investigating if life stress interacts with ELS to cause reward processing deficits. Participants with a history of ELS showed evidence for decreased positive feedback sensitivity in the PRLT, however neither high ELS nor control participants showed a response bias in the points based PRT. As this was the first reported attempt to utilise the PRT in an online environment and due to the failure of both groups to meet the primary endpoint we recruited an additional population of control participants using a modified task design using direct compensation to validate that the PRT can be successfully performed online.

By understanding the links between ELS and reward processing deficits as a hypothesised intermediate phenotype in depression this study aims to provide insights into how a person with a history of ELS is rendered at higher risk for depression.

## 2. Methods

All procedures were approved by the Faculty of Life Sciences and Faculty of Science Research Ethics Committee at the University of Bristol and the study protocol was pre-registered (www.osf.io/538yk). All methods were performed in accordance with the declaration of Helsinki in addition to all other institutional and national guidelines. All participants provided full written consent for the collection, analysis and publication of their data which is available open access (www.ofs.io/63e8j) and were reimbursed at a rate of £6.00 per hour.

### 2.1 Participants

A total of 586 participants were recruited using the Prolific (www.prolific.co) online platform to complete an online screening questionnaire (see supplementary Fig 1 for study overview and table 1 for participant demographics). These participants were 25 - 65 years of age, fluent in English, resident in the UK and had no mild cognitive impairments or dementia. Participants completed the early life stress questionnaire [29] (ELSQ) which asks if participants had prior exposure to specific adverse childhood experiences (ACEs). Participants were also asked “Have you got a current diagnosis of a mental health disorder or Parkinson’s disease?”.

**Table 1.**
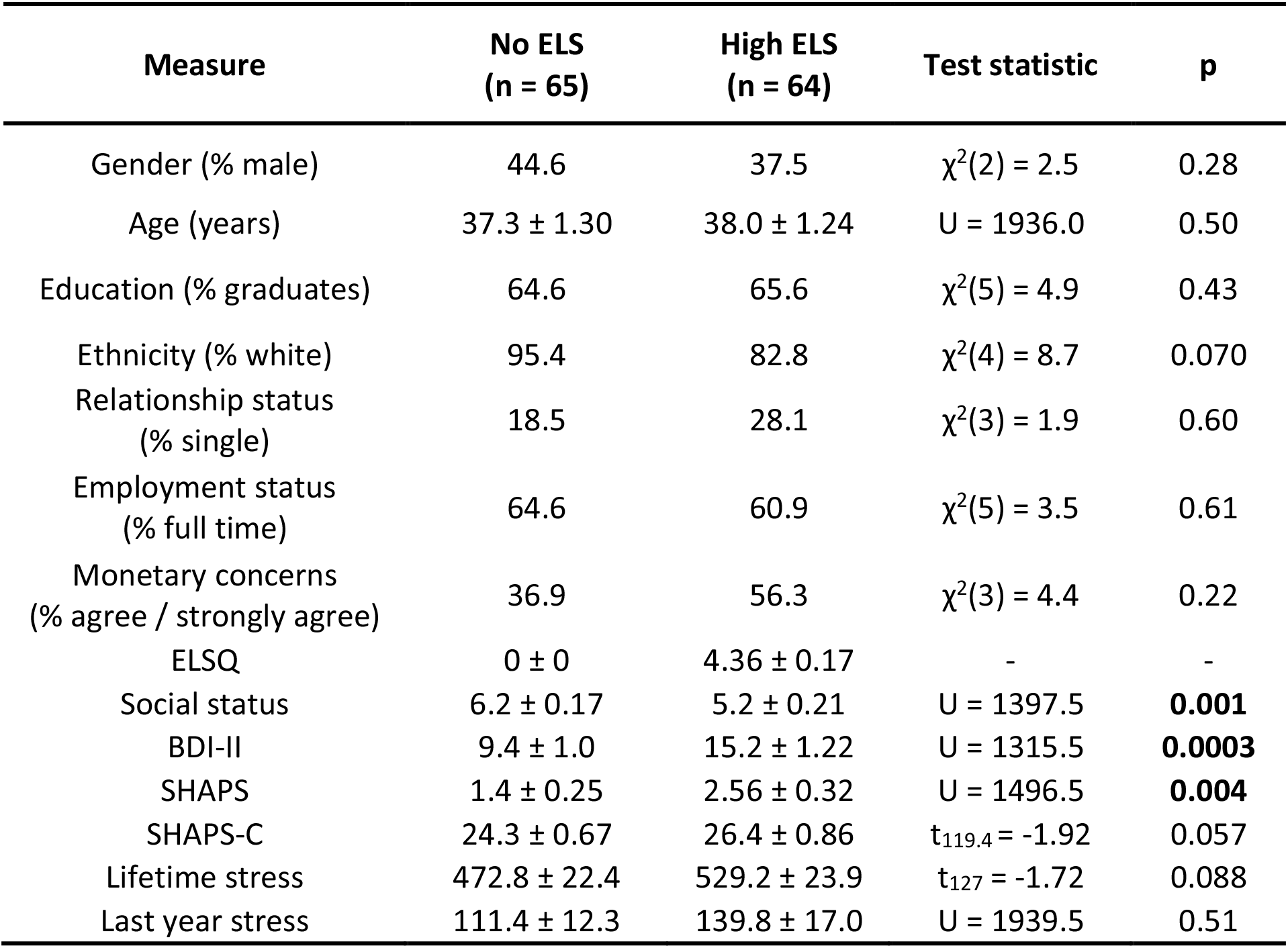
Demographic and self-report measures in the study population. Values are shown for each group as mean ± standard error with significant p values (p ≤ 0.05) indicated in bold.

| Measure | No ELS<br>(n = 65) | High ELS<br>(n = 64) | Test statistic | p |
| --- | --- | --- | --- | --- |
| Gender (% male) | 44.6 | 37.5 | $\chi^2(2) = 2.5$ | 0.28 |
| Age (years) | 37.3 ± 1.30 | 38.0 ± 1.24 | U = 1936.0 | 0.50 |
| Education (% graduates) | 64.6 | 65.6 | $\chi^2(5) = 4.9$ | 0.43 |
| Ethnicity (% white) | 95.4 | 82.8 | $\chi^2(4) = 8.7$ | 0.070 |
| Relationship status<br>(% single) | 18.5 | 28.1 | $\chi^2(3) = 1.9$ | 0.60 |
| Employment status<br>(% full time) | 64.6 | 60.9 | $\chi^2(5) = 3.5$ | 0.61 |
| Monetary concerns<br>(% agree / strongly agree) | 36.9 | 56.3 | $\chi^2(3) = 4.4$ | 0.22 |
| ELSQ | 0 ± 0 | 4.36 ± 0.17 | - | - |
| Social status | 6.2 ± 0.17 | 5.2 ± 0.21 | U = 1397.5 | **0.001** |
| BDI-II | 9.4 ± 1.0 | 15.2 ± 1.22 | U = 1315.5 | **0.0003** |
| SHAPS | 1.4 ± 0.25 | 2.56 ± 0.32 | U = 1496.5 | **0.004** |
| SHAPS-C | 24.3 ± 0.67 | 26.4 ± 0.86 | $t_{119.4} = -1.92$ | 0.057 |
| Lifetime stress | 472.8 ± 22.4 | 529.2 ± 23.9 | $t_{127} = -1.72$ | 0.088 |
| Last year stress | 111.4 ± 12.3 | 139.8 ± 17.0 | U = 1939.5 | 0.51 |

Participants who met the inclusion criteria for high ELS or no ELS and did not report a diagnosis of a mental health disorder or Parkinson’s were then invited to take part in a second phase of the experiment online within a week of screening and were allocated into two groups. A no ELS group (n = 65) contained people reporting no ACEs on the ELSQ while a high ELS group (n = 64) consisted of those who reported ≥3 ACEs (estimated to be the top tercile of the population [29]). In this second phase of the experiment participants entered demographic information before completing the MacArthur Scale of Subjective Social Status [30], Beck’s depression inventory II [31] (BDI-II), the Snaith Hamilton pleasure scale [32] (SHAPS) and the Holmes and Rahe stress scale [33]. The SHAPS was additionally scored using the SHAPS-C criteria [34]. For the stress scale participants were asked if each event occurred in either their adult life or the last year to provide a measure of both total adult lifetime stress and recent stress. For all stages of the experiment participants were instructed to use a desktop or laptop only and that they should be in a quiet place with minimal distractions. Sample size was estimated for a medium effect size (Cohen’s *d* = 0.5) and 80% power for a t-test at 64 participants per group. While other studies have investigated different dimensions of ELS (i.e. emotional abuse vs psychosocial neglect) with regards to cognitive outcome [8] the present study was not powered to enable this.

Early life stress was highly prevalent in the study population with only 21.0% of participants having no adverse childhood experiences (ACEs) and 44.4% of the population suffering three or more ACEs in their childhood (see supplementary fig 2). 16.0% of respondents self-reported a diagnosis of a mental health disorder or Parkinson’s with this being associated with a higher ELSQ score (Mann-Whitney, U = 15725, p < 0.0001).

The two study groups were well matched with respect to gender, age, education, ethnicity, relationship status, employment status and the presence of monetary worries (see table 1). However, high ELS participants had a self-reported lower social status coupled with higher depression scores in the BDI-II and elevated anhedonia scores in the SHAPS questionnaires. There was no evidence of a difference between groups when participants were asked about stress they encountered in both the last year and their adult lives. When the BDI-II scores were classified into either minimal, mild, moderate or severe depression (see supplementary fig 3 [31]) participants from the high ELS group were more likely to be in greater severity depression groupings (chi^2^, χ^2^(3) = 12.9, p = 0.005). Similarly when SHAPS scores were classified into either normal (≤2) or abnormal (≥3) hedonic responses [32] members of the high ELS group were more likely to have abnormal scores (see supplementary fig 3, chi^2^, χ^2^(1) = 6.3, p = 0.012).

### 2.2 Behavioural testing

Following completion of self-report measures, participants completed the Probabilistic reversal learning task [35,36] followed by the Probabilistic reward task [37]. To complete the tasks participants were required to download and install the Millisecond Inquisit web player (Millisecond, US) which ran both tasks using Millisecond Inquisit v6.2.1. Participants were instructed they were able to earn an additional £2.00 for high performance on the behavioural tasks.

#### 2.2.1 Probabilistic Reversal Learning task

The PRLT was conducted as previously described [35,36] using the task from the Millisecond test library [38]. Participants were instructed to choose between a “lucky” (rich) and “unlucky” (lean) pattern to maximise points. Selection of the rich stimulus enabled participants to gain a point 80% of the time and lose a point 20% of the time with the lean stimulus having opposite contingencies. If no stimulus was chosen within 2s then this was classified as incorrect and participants lost a point. After meeting the reversal criterion, the contingencies reverse such that the rich stimuli becomes lean and vice versa. This criterion was set randomly between 10 to 15 consecutive correct rich choices to stop participants counting to the criterion. Participants first completed a practice phase where they had to achieve the criterion for a single reversal before proceeding to the main task which was completed in three blocks each limited to 9 minutes where participants could reverse as many times as able within a block. Participants who did not pass the practice phase were excluded from analysis. Data were analysed as previously described [39]. Win-stay probability was defined as the probability that if a participant was rewarded for selecting a stimulus they would select the same stimulus for the next trial. Lose-shift probability was conversely the probability that if a participant lost a point at a stimulus they would switch to the opposite stimulus for the next trial. This enabled win-stay and lose-shift probabilities to be used as measures of positive and negative feedback sensitivity respectively. These were subdivided into either true, feedback that matches with the underlying task rules (e.g. being rewarded for selecting the rich stimulus), or misleading feedback (e.g. being rewarded for selecting the lean stimulus), that which is opposite to the underlying task rule. The number of rule changes, how many times participants were able to meet criterion for a rule change, accuracy and response latency per block were additionally analysed. A Qlearn reinforcement learning model was applied to data as previously described [39,40] to give estimates of model free learning rate, accuracy compared to a model predicted perfect strategy (subjective accuracy) and beta, a measure of choice variability (low values indicate essentially random choices while high values show a deterministic choice strategy). Additionally, data per phase (practice, acquisition of the first rule in block 1 and the following two reversals) was analysed consisting of participant accuracy, errors to criterion and win-stay / lose-shift probability.

#### 2.2.2 Probabilistic Reward Task

The PRT was conducted as previously described [37] using the task from the Millisecond test library [41]. Participants were instructed to identify whether the mouth of a presented cartoon face was long or short (≈11% difference in mouth length) to win points over 3 blocks of 100 trials. Participants were shown a face before a mouth was rapidly presented for 100ms with participants given up to 1750ms to respond. Feedback was not provided on every trial but unknown to participants one mouth was rewarded with points three times more often than the other (rich = 60%, lean = 20%). Response key and rich/lean stimuli assignments were counterbalanced across participants and responses that were quicker than 150ms or slower than 1750ms were excluded from analysis. Additional responses that differed by more than 3 standard deviations from the mean following natural log transformation of latencies for each participant were excluded from analysis. Response bias (logB), a measure of reward learning, and discriminability (logD), a measure of task difficulty, were calculated as:

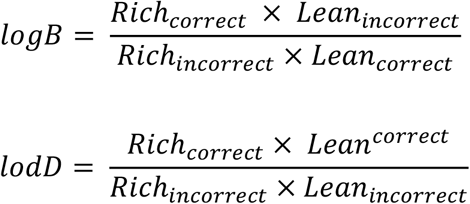

#### 2.2.3 Directly rewarded probabilistic reward task

An additional cohort of 81 participants (no current or previously diagnosed mental health condition, aged 18-45) were recruited using the Prolific platform (see supplementary methods for more details). Participants completed a different reward learning task (not discussed here) over 5 days and on the final day completed the directly rewarded PRT alongside BDI and SHAPS questionnaires. The directly rewarded PRT was carried out identically to as previously described except participants were given direct monetary compensation for each rewarded trial (£0.04) as opposed to points which later converted to a bonus payment. Only participants with a minimal BDI score (<13) and normal SHAPS score (≤2) were included in final analysis.

### 2.3 Data Analysis

Demographic and self-report measures were compared between groups using either χ^2^, t-tests or Mann-Whitney U tests where appropriate. The primary analysis for each measure was a direct comparison between no ELS and high ELS groups. Where data were not normally distributed then efforts were first made to transform data to normality and where this was not possible Mann-Whitney U tests were completed. Win-stay by block data were transformed using the bestNormalize package in R [42]. Where measures were split by a within subject factor such as block or feedback type these were analysed with repeated measures ANOVAs. Where Mauchly’s test identified a violation of the Sphericity assumption then this was corrected using the Huynh-Feldt correction. T-tests were used for direct group comparisons.

Due to differences in social status, BDI-II score and SHAPS score between the no ELS and high ELS groups, principal component analysis (PCA) was conducted to reduce the dimensionality of these variables to account for depression symptomology as an analysis stage (see supplementary tables 1 and 2). Because only principal component 1 (PC1) differed between groups and explained 94.6% of variance this was used in ANCOVAs (analysis of covariance) to analyse whether parameter changes were due to ELS or due to changes in depression symptomology accounted for by the PC1 component. To understand if stress and gender interacted with ELS to modify reward learning, exploratory analysis was also undertaken using generalised linear mixed models (GLMMs) containing the factors: gender, ELS, lifetime stress, last year stress and age. GLMMs were fit using the glmmTMB package in R 4.0 [43,44] with model refinement conducted utilising stepwise deletion based upon Akaike information criterion before being compared with a null model to protect against overfitting. PC1 was also added to each model following final model selection to assess the effects of depression symptomology.

Statistical analysis was conducted in SPSS v26 (IBM, US), MATLAB 2018a (Mathworks, USA) and R 4.0 [43] with output graphics constructed in GraphPad Prism 8 (GraphPad, US). For all analysis α was set at 0.05. All data is shown as mean ± SE with a bar and stars showing evidence of a main effect of ELS in the primary analysis. * ≤ 0.05, ** < 0.01, *** < 0.001, **** < 0.0001.

## 3. Results

### 3.1 Probabilistic reversal learning task

There was no evidence of a difference between groups in either the number of rule changes participants were able to complete (Fig 1A) nor accuracy (Fig 1B). However, participants with a history of high ELS did have a slower average response latency (Fig 1C, RM-ANOVA, F_1,126_ = 5.03, p = 0.027) with both groups getting equally faster over the course of the three blocks (RM-ANOVA, F _1.88,236.7_ = 16.1, p < 0.0001). Secondary analysis revealed no evidence of an effect of depression symptomology (RM-ANCOVA, PCA1: p > 0.05) with the main effect of ELS persisting (RM-ANCOVA, ELS: F_1,125_ = 4.9, p = 0.028). Exploratory analysis on overall reaction times did not replicate a main effect of group but did observe older participants having slower reaction times (GLMM, Z = 2.8, p = 0.005). This analysis also indicated weak evidence of an interaction between group and lifetime stress (GLMM, Z = 1.55, p = 0.065) but further investigation did not reveal an effect of lifetime stress in either group.

**Fig 1.**
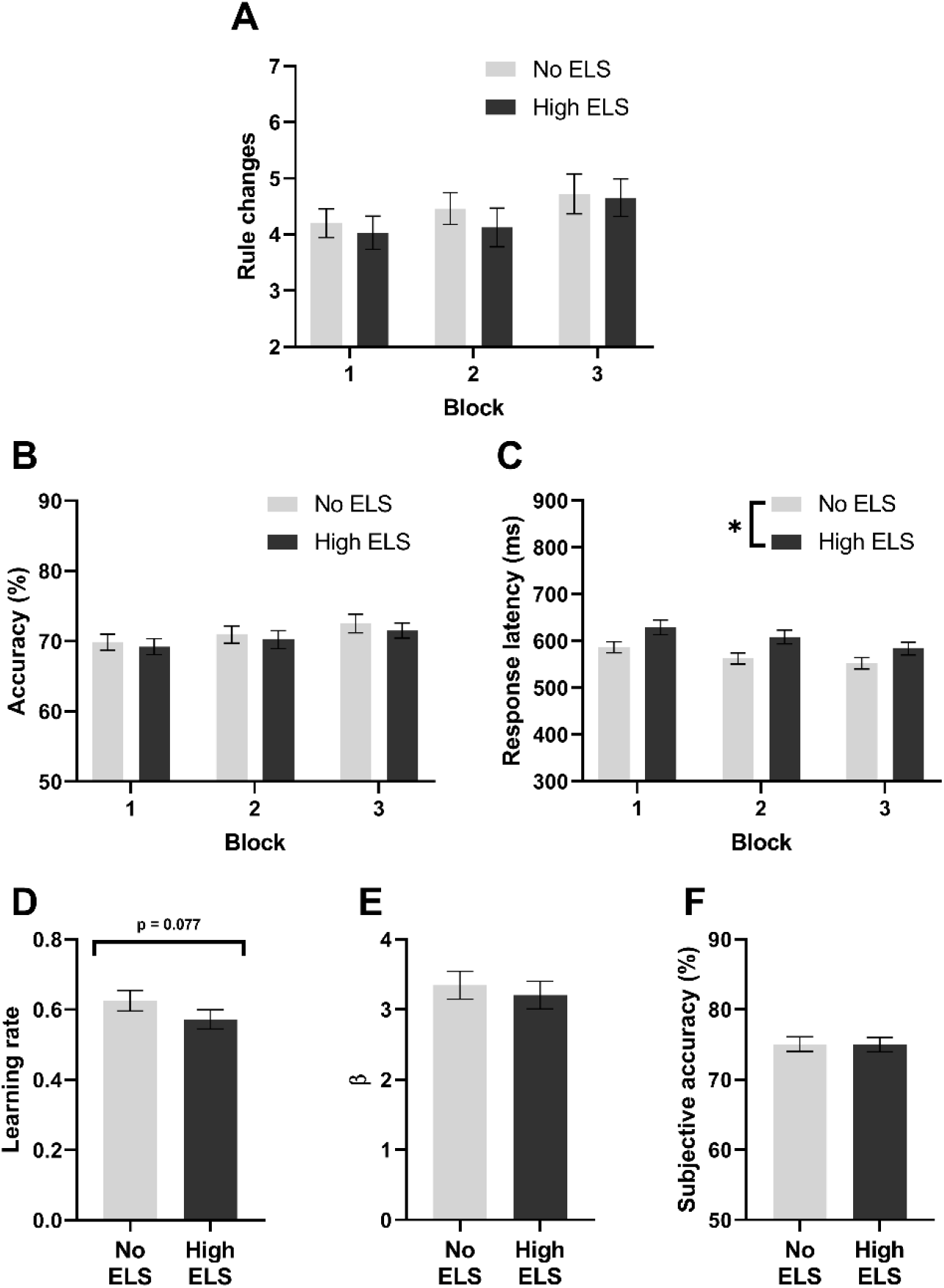
Overall reward learning and reinforcement learning in the PRLT. **(A)** Rule changes within each block, **(B)** accuracy by block and **(C)** average response latency per block. From the Q-learning reinforcement learning model: (**D)** learning rate, **(E)** β, the inverse of the softmax temperature and a measure of choice variability and **(F)** subjective accuracy, participant accuracy compared to a model predicted perfect strategy.

When data were analysed using the Q-learning reinforcement learning model a trend emerged towards high ELS participants having a lower learning rate compared to the no ELS study population (Fig 1D, t-test, t_127_ = 1.78, p = 0.077). Secondary analysis revealed no effect of PCA component 1 upon learning rate but removed any evidence for an effect of ELS. In exploratory analysis a main effect of ELS was observed (GLMM, Z = 2.1, p = 0.037) with the addition of PC1 impairing model fit (ΔAIC = 1.69, χ^2^(1) = 0.31, p = 0.57). Additionally, a relationship between stress in the last year and learning rate was observed whereby increased stress in the last year decreased learning rate (GLMM, Z = −2.3, p = 0.024). There was no difference in choice variability (Fig 1E) or accuracy compared to a model predicted perfect strategy (Fig 1F) between groups.

Participants with a history of high ELS exhibited reduced positive feedback sensitivity (PFS, Fig 2A, RM-ANOVA, F_1,122_ = 10.4, p = 0.002) which persisted once depression symptomology was accounted for using PCA component 1 (RM-ANOVA, F_1,121_ = 6.6, p = 0.01). Exploratory analysis revealed an interaction between ELS and both lifetime stress (GLMM, Z = −2.15, p = 0.031) and last year stress (GLMM, Z = −1.99, p = 0.047). Further investigation revealed effects of both stress types upon PFS in the low ELS group only (GLMM, lifetime stress: Z = −2.35, p = 0.019, last year stress: Z = −2.2, p = 0.026) whereby higher lifetime stress led to greater PFS but higher stress in the last year was associated with decreased PFS. However it should be noted that although all suggested terms were removed from the model the overall model was a poorer fit than the null when measured by AIC (Δ AIC = 7.3, χ^2^(13) = 18.7, p = 0.13).

**Fig 2.**
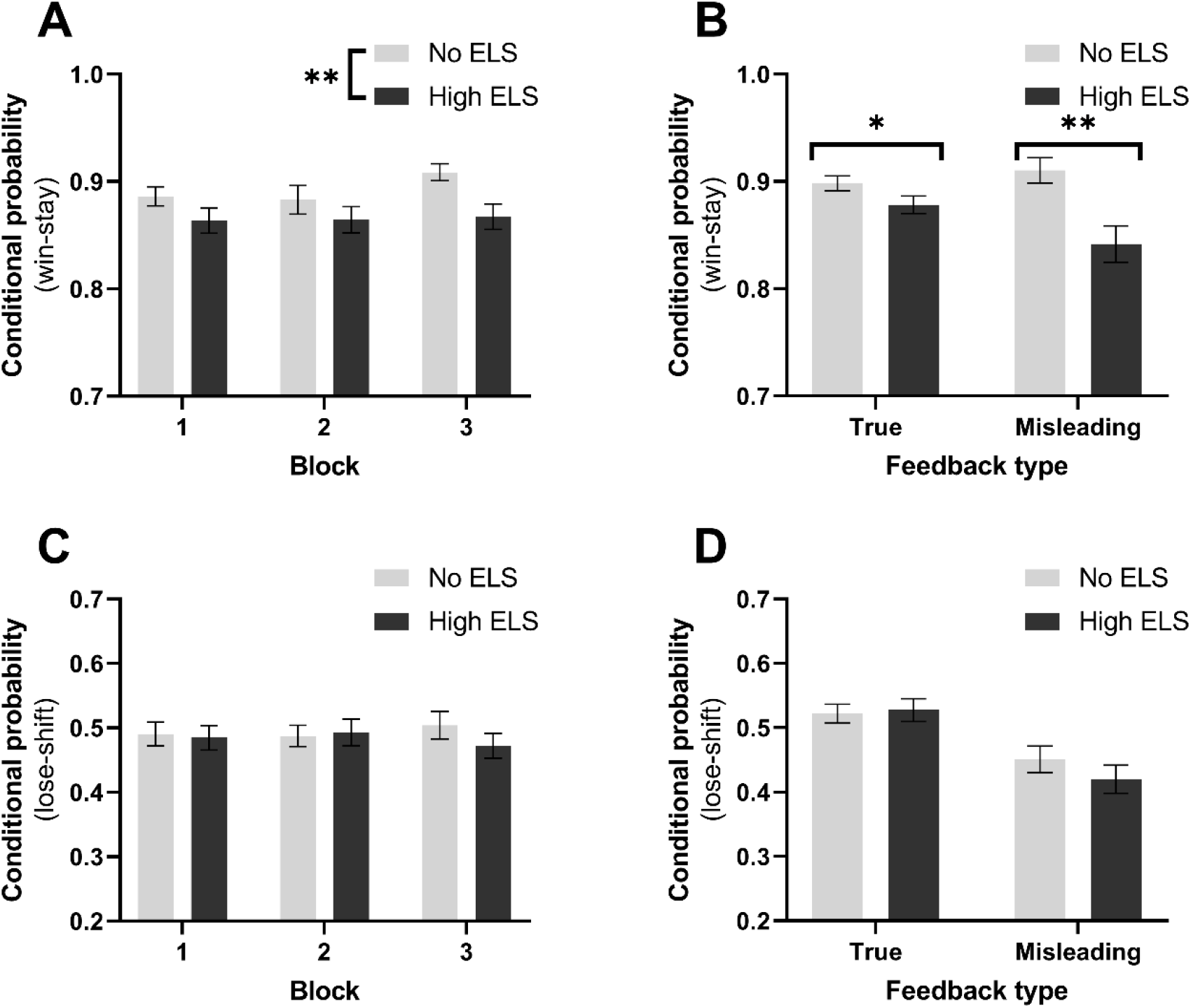
High ELS participants exhibited lower positive feedback sensitivity than those without a history of ELS. Win-stay probability overall **(A)** and subdivided into true and misleading feedback **(B)**. Overall Lose-shift probability **(C)** and additionally subdivided into true and misleading feedback (**D)**.

The effect of ELS upon PFS was consistent across feedback that matched (true feedback) or clashed (misleading feedback) with the underlying task rules (Fig 2B, Mann-Whitney U, true: U = 1443, p = 0.03; misleading: U = 1337, p = 0.005). This effect appeared to be constrained to PFS with no corresponding changes in lose-shift probability between no ELS and high ELS groups (Figs 2C and D).

When initial learning in the PRLT task was assessed, it was apparent that although ELS and control participants performed similarly during the practice phase there was a learning deficit during acquisition of the first reversal criterion in block 1 as evidenced by increased errors to criterion (Fig 3A, Mann-Whitney U, U = 1580, p = 0.045) and decreased accuracy (Fig 3B, Mann-Whitney U, U = 1584, p = 0.036). Both groups of participants however performed equally well at achieving criterion for a second and third reversal. Unlike the overall measures there was no difference in win-stay probability between groups (Fig 3C), however there was a trend for high ELS participants to show increased negative feedback sensitivity (NFS) in the practice phase (Fig 3D, Mann-Whitney U, U = 1532, p = 0.052).

**Fig 3.**
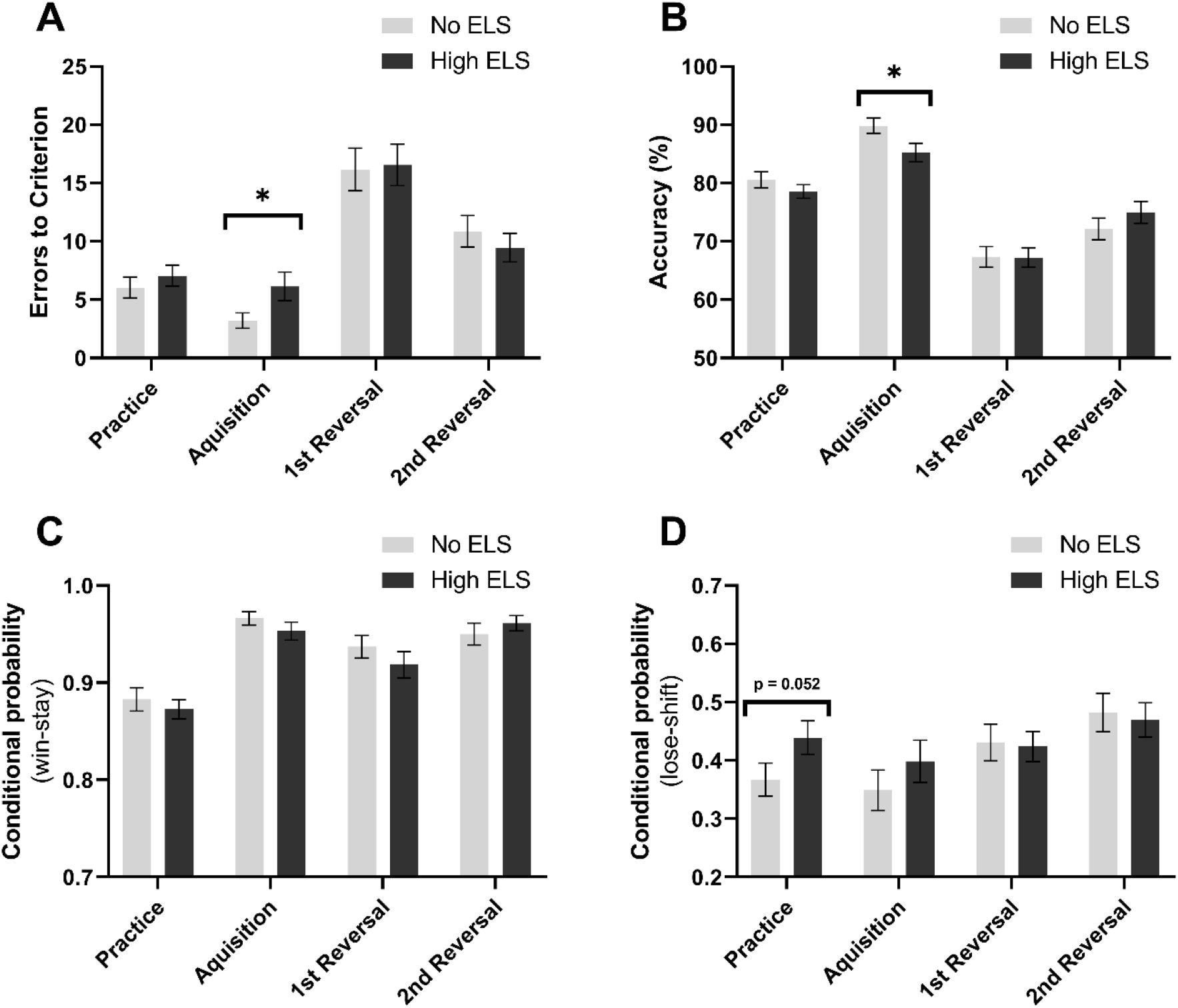
High ELS participants show impaired learning in the acquisition phase of block 1. **(A)** Errors made while reaching criterion for each phase, **(B)** accuracy within each phase, **(C** and **D)** win-stay and lose-shift probabilities for each phase of block 1 and practice respectively.

### 3.2 Probabilistic reward task

There was no evidence that participants developed a response bias towards the more highly rewarded stimulus in any block (Fig 4A) nor was there evidence for a response bias developing between blocks (Fig 4B). However, participants with a history of high ELS did show an impaired ability to discriminate between stimuli (Fig 4C, ANOVA, F_1,127_ = 4.8, p = 0.030). Secondary analysis revealed that this difference between groups appeared to be driven by differences in depression symptomology with the effect of ELS disappearing when PCA component 1 was included in the analysis (ANCOVA, PCA1: F_1,126_ = 6.08, p = 0.015; ELS: F_1,126_ = 1.7, p = 0.19). Exploratory analysis further revealed a main effect of lifetime stress with higher lifetime stress corresponding to increased discrimination ability (GLMM, Z = 2.6, p = 0.007). An effect of gender was also revealed (GLMM, Z = 2.04, p = 0.04) with males showing increased discrimination ability. Finally, there was no difference between groups in response latencies (Fig 4D) nor was there an effect of stimulus upon response latency (Fig 4E).

**Fig 4.**
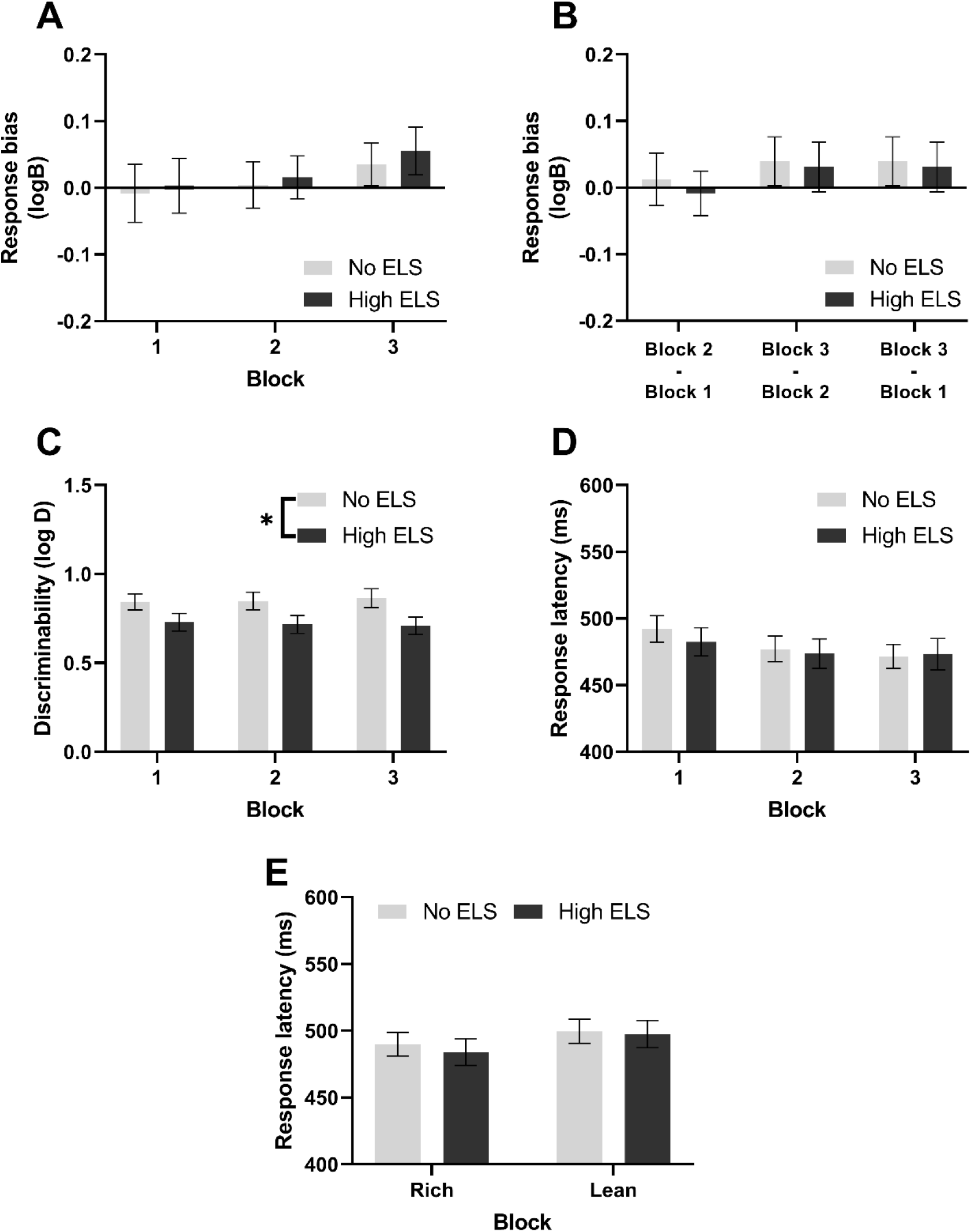
Participants with a history of ELS show decreased discriminability in the PRT. **(A)** Response bias to the more highly rewarded stimulus, **(B)** response bias development between blocks, **(C)** discriminability between long and short face lengths, **(D)** average response latency split by block and **(E)** response latency split by stimulus type.

Consistent with Pizzagalli and colleagues [19] the probability of misclassifying a stimulus based upon the preceding trial outcome was also analysed (supplementary table 3). Participants with a history of high levels of ELS were more likely to misclassify rich stimuli if either the previous trial was a not rewarded rich trial or a lean not rewarded trial with these measures roughly corresponding with rich lose-shift and lean lose-stay probability in the PRLT respectively.

### 3.3 Directly rewarded probabilistic reward task

To further investigate the lack of overall response bias in the online version of this task, a second cohort of control participants completed the PRT using direct monetary compensation (£0.04 reward per correct trial) instead of points reward (see supplementary materials). Participants showed a response bias in blocks 1 and 3 (Fig S4A, Wilcoxon signed ranks test, block 1: W = 1087.5, p = 0.001, block 3: W = 916.5, p = 0.038) but there was no overall effect of block. There was no evidence for response bias increasing over time (Fig S4B) nor was there evidence for any effect of block upon discriminability (S4C) and response latency (S4D). Participants did however respond more quickly to rich stimuli than lean (Fig S4E, Wilcoxon matched pairs signed ranks test, W = 814.0, p = 0.0007).

## 4. Discussion

This study was designed to investigate whether healthy adults with a history of ELS show alterations in reward processing and feedback sensitivity. Nearly 600 participants were screened; ELS was highly prevalent in the population with 79.0% of participants experiencing one or more ACE and 44.4% experiencing three or more.

Participants with a history of high ELS had higher self-report depression and anhedonia symptoms. Although participants stated they did not have a diagnosis of depression, 54.7% of high ELS and 26.2% of no ELS participants showed at least mild symptoms based upon the BDI-II questionnaire. BDI-II scores in no ELS (mean: 9.4 ± 1.0) and high ELS (mean: 15.2 ± 1.2) participants were higher than controls for similar studies [19,28,45] (range 1.3-3.62) but lower than depressed patients [19] (mean: 32.1 ± 8.6) or participants described as exhibiting a high BDI [45] (>16). These data are consistent with a large societal burden of un-diagnosed depression [46–48]. It should be noted that the present study was undertaken during the Covid-19 global pandemic with it being estimated that levels of depression had doubled during this period [49]. The high number of participants reporting mild to severe symptoms of depression is a major limitation of this study as depression and anhedonia are well known to reduce reward learning in both the PRLT [24] and PRT [18,19,37]. It is also worth considering that 75% of adults with mental health conditions experience the onset of symptoms before aged 24 [50]. This means that the study population, all 25 years of age or greater, is potentially biased towards those more protected from mental health disorders. In keeping with large bodies of previous literature this study used retrospective reporting however it is worth noting the difficulties in correlating retrospective and prospective measures of ELS [51] suggesting that these might encompass different populations with different mechanistic links between ELS and depression vulnerability. Studies have also interesting insights when ELS is split into different modalities [52,53], however the present study was not powered enough to merit investigating this.

### 4.1 Probabilistic reversal learning task

In the PRLT participants with high ELS displayed decreased positive feedback sensitivity compared to controls as measured by win-stay probability. This finding was independent of depression symptomology and specific to PFS with no changes observed in lose-shift probability. Blunted striatal responses to reward in participants with a history of ELS have been previously reported [54,55] which speculatively may underlie the decreased PFS observed in the present study. Consistent with the present study, women with MDD and a history of childhood sexual abuse have also been found to have impaired performance in the PSST but only for trials requiring use of previously rewarded information and not those requiring use of previously punished information [28]. Within the PRLT depressed patients have been observed to show increased sensitivity to misleading negative feedback [23,24]. This was not observed in the high ELS cohort in the present study. In other tasks depressed patients have also been reported to show increased NFS alongside attenuated PFS [56–60]. These findings suggest that ELS is associated with changes in feedback sensitivity in the PRLT differently to depression with ELS decreasing PFS but not effecting NFS while depression has an opposite effect.

The PRLT also allows for assessment of reinforcement learning through the analysis of rule changes and accuracy in addition to parameters calculated through use of the reinforcement learning model. In contrast with our hypothesis, there was no evidence that ELS affected rule changes which is surprising considering evidence that both depression and ELS can impair cognitive flexibility [61,62]. Although rule changes were used as the main behavioural reward learning output, when data were analysed with the Q-learning model a trend towards decreased learning rate was observed in high ELS participants. This became significant in exploratory analysis and decreased associative learning has been previously observed in juveniles previously exposed to physical abuse [27]. There is a lack of consistent evidence in depression studies as to whether model free learning rate differs between patients and controls [63,64]. These findings warrant future investigation due to this study being only powered to detect group differences between two groups meaning that ANCOVA and exploratory analysis is likely to be underpowered.

A slower response latency was also observed in high ELS participants which was specific to the PRLT with no congruent changes seen in the PRT. This discrepancy may be related to differing cognitive demands with the PRLT potentially requiring greater working memory.

No directly comparable studies have been carried out in humans. However, maternally separated marmosets, an animal model of ELS, showed no change in simple visual discrimination compared to controls but showed impairments when the contingencies reversed [65]. This is similar to that seen in both depressed and bipolar patients in the human PRLT [24,66] who acquire the initial rule successfully but then are impaired following reversal. This compares to ELS participants in the present study who performed equally well in the practice phase and reversal phases but showed a deficit in acquisition of the first rule in block 1. This suggests a potential impairment in the ability to generalise the task rules between the practice and acquisition phase. However previous probabilistic learning studies did not include a practice phase meaning that this likely changed the way participants processed the start of block 1. This might explain the contrast with Pechtel and Pizzagalli, 2013 who reported that women with remitted MDD and ELS learnt acquisition in the PSST at the same rate as controls [28].

One of the hypotheses of this study was that stress in adult life would modulate the relationship between reward processing deficits and ELS. There was little evidence that this was the case except for an observed interaction between PFS and ELS whereby stress only influenced PFS in participants without a history of ELS. Higher lifetime stress led to greater PFS but higher stress in the last year was associated with decreased win-stay probability. There are few previous studies investigating similar constructs but Berghorst and colleagues reported that after stress induction those who had higher cortisol reactivity and self-reported negative affect had lower reward but not punishment sensitivity [26]. Additionally, it is worth noting that due to the relatively poor model fit for this exploratory analysis that these findings should be taken as preliminary due to the risk of data overfitting.

### 4.2 Probabilistic reward task

In contrast to previous studies employing the PRT neither ELS nor control groups showed a response bias toward the more highly rewarded stimulus [19,37] suggesting a general failure of all participants to modulate their responses as a function of reward. This lack of response bias in the main study potentially indicates that the reward information was not salient enough therefore participants focussed upon correctly discriminating between mouth lengths. This makes comparison to previous literature challenging. There are no previously published studies carrying out the PRT online but in this study we failed to replicate the main outcome measure. The online testing could have been one reason for the lack of response bias likely leads to the high variability seen in the data as it is not possible to ensure that participants are completing the tasks in as controlled an environment as would be possible by laboratory testing.

However, another possibility for the lack of response bias seen was a key difference between laboratory and online versions of the task. While all other aspects of the task were similar, participants in the online task were informed that high performance would lead to a bonus payment with the actual reward in the task being points. Previous studies instead used direct monetary compensation in the task [37]. When the second population of control participants was tested in the PRT using direct monetary compensation a response bias was seen in blocks 1 and 3, however there did not appear to be evidence for this bias strengthening over time like previously observed [37]. There was also robust evidence for participants responding more quickly to the rich than lean stimulus as also has been reported [22,37]. While this difference in compensatory mechanism may underly the difference in control population performance in the two implementations of the task it could also be because the direct reward population had lower BDI and SHAPS scores. However, when the direct reward experiment was re-analysed with no BDI and SHAPS cut-offs such that it more closely approximated the control population in the main study the results were much the same as with the cut-offs (data not shown). These data therefore suggest that it is possible to successfully implement the PRT in an online setting using the directly rewarded task with the availability of reliable online psychological tasks being key under current circumstances.

While difficult to interpret for previously discussed reasons, participants with high levels of ELS did show impairments in discrimination, a measure of task difficulty. This however appeared to be driven by changes in depression symptomology as opposed to ELS specifically.

### 4.3 Conclusions

These data suggest that participants who do not self-report a diagnosis of a mental health condition but do have a history of ELS show impairments in positive feedback sensitivity and reward learning in the PRLT compared to controls. These impairments may be important in understanding how ELS predisposes to depression with reduced reward learning being a key feature in MDD patients [20]. However, high levels of potentially undiagnosed depression are a potential confound and highlight a potential wider issue in terms of the number of people who meet criteria for MDD but are not formally diagnosed or receiving care. Future studies are needed to replicate these findings, investigate the neural circuit changes underlying these reward learning impairments and investigate whether these findings are directly related to psychiatric risk.

5.

## Acknowledgements

The authors would like to thank Michelle Taylor for advice on statistical analysis. This work was primarily funded by the BBSRC SWBio DTP PhD programme (grant numbers: BB/J014400/1 and BB/M009122/1) awarded to MPW. Additional support was also provided from the Wellcome Trust Neural Dynamics PhD studentship (grant number: 108899/B/15/Z) awarded to CLS. For the purpose of Open Access, the author has applied a CC BY public copyright licence to any Author Accepted Manuscript version arising from this submission.

## 6. Data Availability Statement

This study was pre-registered and available at www.osf.io/538yk. Data and code will also be made available open access at www.osf.io/63e8j.

## Notes

https://osf.io/538yk

https://osf.io/63e8j/

### Competing Interest Statement

The authors have declared no competing interest.

### Summary of Updates

We have additionally included some control experiments using a modified online probabilistic reward task (PRT) in a healthy adult population. Within this experiment we observed a response bias suggesting the online PRT can be successfully used online.

