## Supplementary Materials for "Investigation of reward learning and feedback sensitivity in non-clinical participants with a history of early life stress"

### <u>Supplementary Material</u>

**Supplementary Methods – Directly rewarded PRT in a control population.**

An additional cohort of 81 participants were recruited to assess the online PRT using direct monetary compensation.

**Participants**

Eligibility Criteria were: aged 18 – 45 years, fluent in English, resident in the UK, normal or corrected-to-normal vision (self-report), no current or previous diagnosed mental health condition (self-report), reported using a macOS or windows 10 operating system, not taken part in previous prolific studies from the same researcher (i.e., a prolific blocklist was employed) and scored > 65% "correct" on a different reward learning task (the reward learning assay, not discussed here).

**Procedure**

All participants completed a different reward learning task (the reward learning assay) over five consecutive days. On the final day, participants also completed the Probabilistic Reward Task (PRT)[1], Snaith-Hamilton Pleasure Scale (SHAPS)[2] and Beck Depression Inventory (BDI; suicide question removed)[3].

**Probabilistic Reward Task**

The PRT (Pizzagalli et al., 2005) available on the Millisecond test library was employed (using Inquisit v6). The only change made to the task was the monetary amount of reward: participants were informed that they could win up to £5 on this task. Specifically, they were informed that if a correct response is rewarded they will earn four pence.

**Analysis**

Only participants with a minimal BDI score (<13) and normal SHAPS score (≤2) were included in final analysis. The output variables logB and logD were calculated as described elsewhere [1]. Data were both analysed across all blocks for a variable using Friedman tests due to the non-normality of data with response bias data also being compared against a hypothetical mean of zero for each block using Wilcoxon signed rank tests.

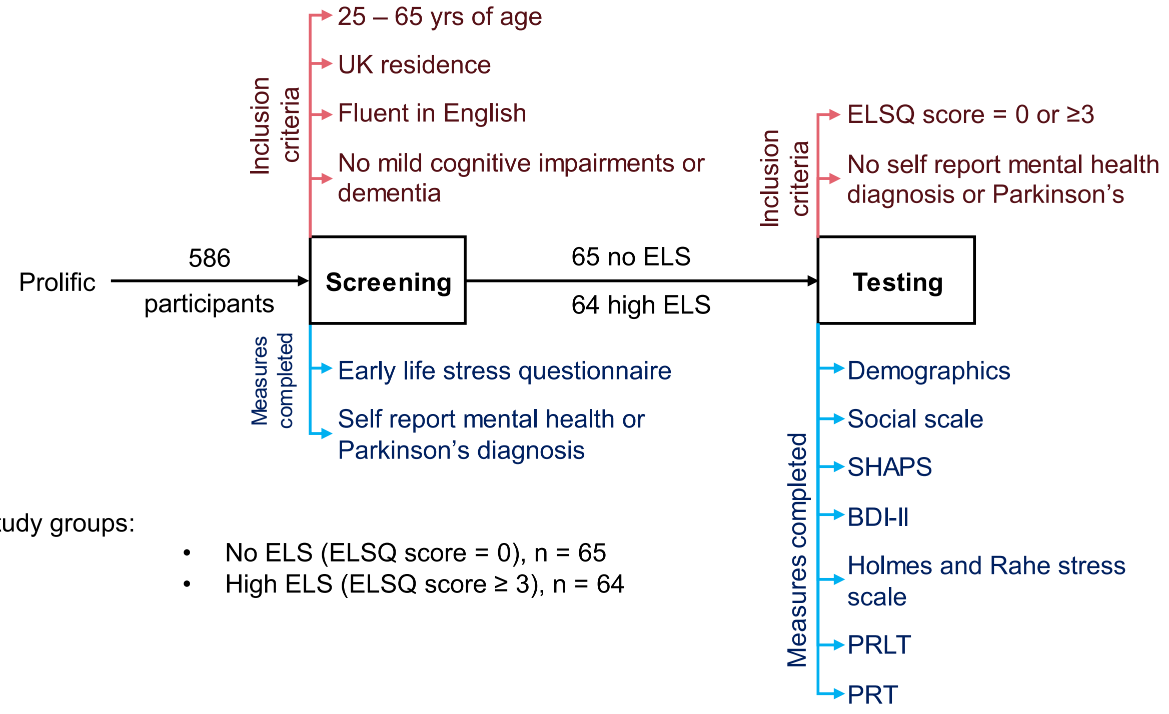


**Fig S1 Study overview.** Participants were screened by ELSQ score and then formed into two study groups: no ELS and high ELS.

| Component | Explained variance (%) | No ELS | High ELS | Test statistic | P value |
|---|---|---|---|---|---|
| 1 | 94.6 | 4.32 ± 0.24 | 5.65 ± 0.25 | $t_{127} = -3.86$ | **0.0002** |
| 2 | 3.4 | -0.19 ± 0.21 | 0.20 ± 0.24 | $t_{127} = -1.22$ | 0.226 |
| 3 | 2.0 | 0.21 ± 0.15 | -0.22 ± 0.18 | $t_{127} = 1.79$ | 0.076 |

**Table S1 Principal component analysis of social scale, SHAPS and BDI-II scores.** The mean ± standard error are shown for each group with the relevant statistical comparison.

| | Principle component | | |
|---|---|---|---|
| | **1** | **2** | **3** |
| Social scale | -0.07 | -0.40 | 0.91 |
| BDI-II | 0.98 | -0.18 | -0.003 |
| SHAPS | 0.17 | 0.90 | 0.40 |

**Table S2 Principal component analysis component loadings.**

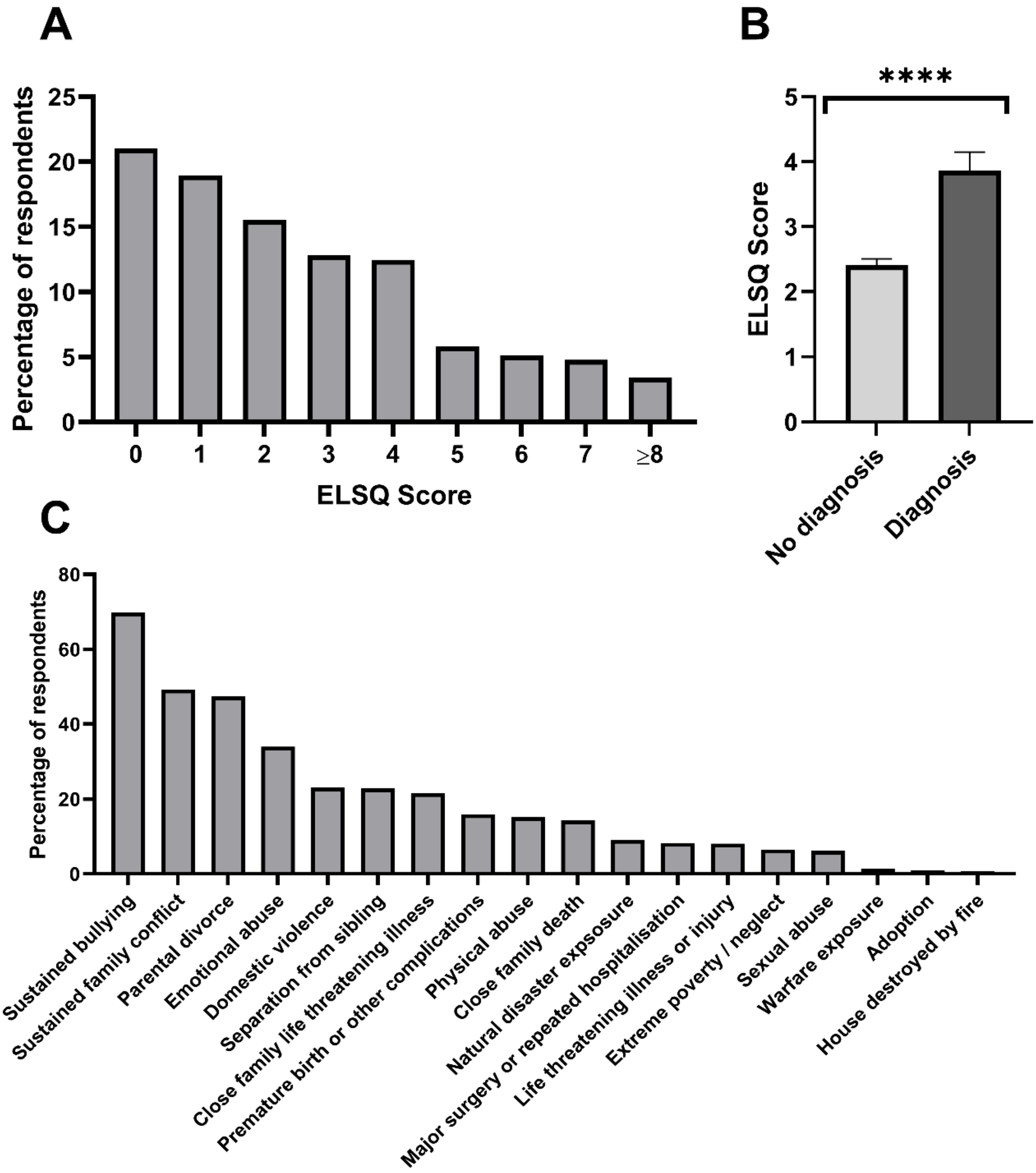


**Fig S2 Early life stress in an online study population. (A)** ELSQ scores in the study population. **(B)** Mental health disorder / Parkinson's self-report diagnosis by ELSQ score (Mann-Whitney, U = 15725, $p < 0.0001$). N = 586 participants. **(C)** ELSQ scores in the study population split by modality of adverse childhood experience.

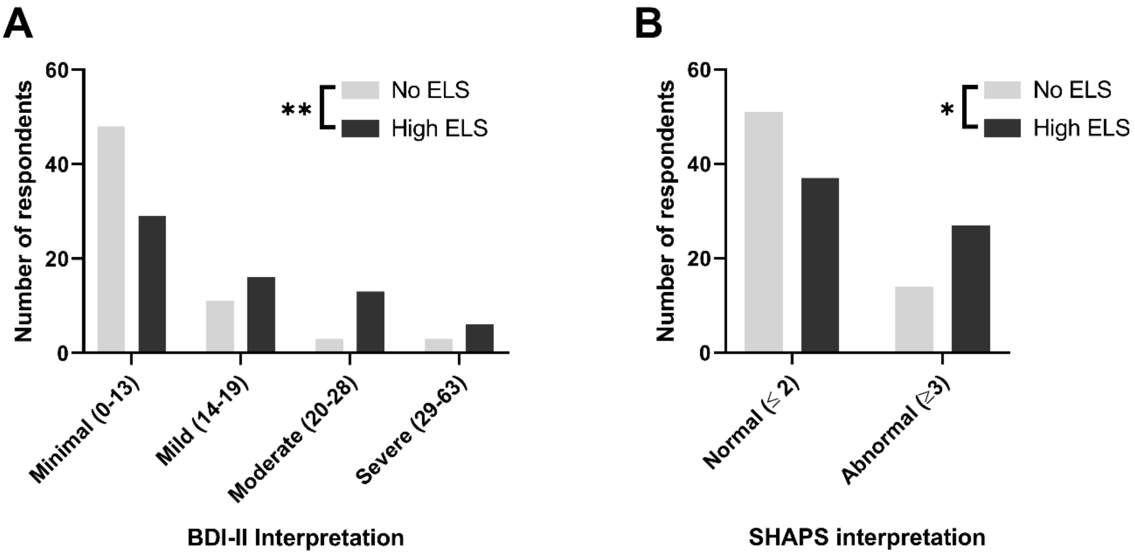


**Fig S3 Interpretation of BDI-II and SHAPS scores in the no and high ELS populations.** Scores were interpreted following Beck et al., 1996 and Snaith et al., 1995. (A) BDI-II split by severity of depression (chi$^2$, $\chi^2(3) = 12.9$, $p = 0.005$) and (B) SHAPS split by normal or abnormal hedonic responses (chi$^2$, $\chi^2(1) = 6.3$, $p = 0.012$). N = 129 participants (65 no ELS, 64 high ELS).

| Current trial | Previous trial | No ELS | High ELS | Test statistic | p |
|---|---|---|---|---|---|
| Lean | Rich - rewarded | 16.8 ± 2.1 | 18.4 ± 2.2 | U = 1966 | 0.59 |
| Lean | Rich - not rewarded | 16.1 ± 1.8 | 18.6 ± 1.8 | U = 1744 | 0.11 |
| Lean | Lean - rewarded | 19.3 ± 2.1 | 20.5 ± 2.0 | U = 1928 | 0.47 |
| Lean | Lean - not rewarded | 16.8 ± 1.6 | 21.2 ± 1.8 | U = 1928 | 0.097 |
| Rich | Rich - rewarded | 13.1 ± 1.5 | 18.3 ± 2.1 | U = 1697.5 | 0.071 |
| Rich | Rich - not rewarded | 14.2 ± 1.6 | 20.0 ± 2.1 | U = 1597.5 | **0.023** |
| Rich | Lean - rewarded | 13.1 ± 1.4 | 15.5 ± 1.7 | U =1814 | 0.330 |
| Rich | Lean - not rewarded | 14.2 ± 1.4 | 19.6 ± 1.9 | U = 1644.5 | **0.040** |

**Table S3 Miss-rates, the chance of mis-categorising a stimulus, by previous trial.** Data is shown as mean ± standard error and significant p-values are shown in bold.

| Measure | Control population (n = 56) |
|---|---|
| Sex (% Male) | 51.8 |
| Age (years) | 31 ± 1.1 |
| Employment (% full time) | 48.2 |
| Student status (% student) | 28.6 |
| BDI | 4.1 ± 0.3 |
| SHAPS | 0.30 ± 0.09 |
| SHAPS-C | 20.7 ± 0.6 |

**Table S4. Demographic and self-report measures in the directly rewarded PRT control population.**

Values are shown for each group as mean ± standard error where appropriate.

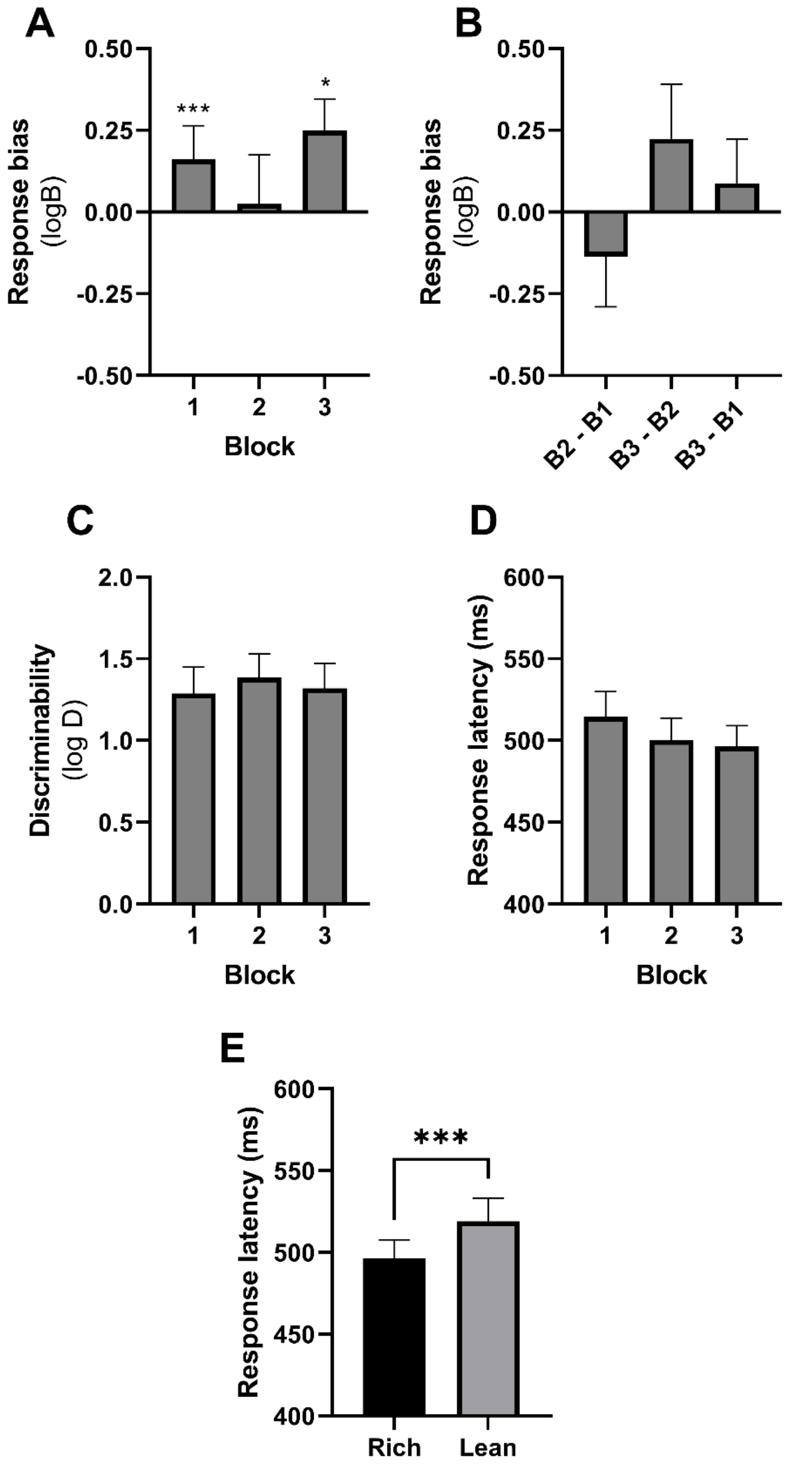


**Fig S4. Direct monetary reward in the PRT using a control population led to a reward induced bias.** **(A)** While no overall effect of block was observed, a response bias was observed in blocks 1 and 3 (Wilcoxon signed ranks test, block 1: W = 1087.5, p = 0.001, block 3: W = 916.5, p = 0.038). **(B)** There was little evidence for response bias strengthening across blocks. Discriminability **(C)** and response latency **(D)** did not appear to change over the course of a session. **(E)** Participants were faster to respond to the rich stimulus than lean (Wilcoxon matched pairs signed ranks test, W = 814, p = 0.0007). N = 56 participants.
